# CLPB3 is required for the removal of chloroplast protein aggregates and for thermotolerance in Chlamydomonas

**DOI:** 10.1101/2022.09.28.509957

**Authors:** Elena Kreis, Justus Niemeyer, Marco Merz, David Scheuring, Michael Schroda

## Abstract

In the cytosol of plant cells, heat-induced protein aggregates are resolved by ClpB/Hsp100 family member HSP101, which is essential for thermotolerance. For chloroplast family member CLPB3 this is less clear with controversial reports on its role in conferring thermotolerance. To shed light onto this issue, we have characterized two *Chlamydomonas reinhardtii clpb3* mutants. We show that chloroplast CLPB3 is required for resolving heat-induced protein aggregates containing stromal TIG1 and the small heat shock proteins HSP22E/F *in vivo* and for conferring thermotolerance under heat stress. Although CLPB3 accumulates to similarly high levels as stromal HSP70B under ambient conditions, we observed no prominent constitutive phenotypes. However, we found decreased accumulation of the ribosomal subunit PRPL1 and increased accumulation of the stromal protease DEG1C in the *clpb3* mutants, suggesting that reduction in chloroplast protein synthesis capacity and increase in protease capacity may compensate for loss of CLPB3 function. Under ambient conditions, CLPB3 was distributed throughout the chloroplast but reorganized into stromal foci upon heat stress, which mostly disappeared during recovery. CLPB3 foci were localized next to signals from HSP22E/F, originating largely to the thylakoid membrane occupied area. This suggests a possible role for CLPB3 in disentangling protein aggregates from the thylakoid membrane system.

**Highlight:** Chloroplast CLPB3 in Chlamydomonas is required for resolving heat-induced protein aggregates and this activity confers thermotolerance under severe heat stress.

During heat stress, CLPB3 organizes into stromal foci located next to the thylakoid membrane system, indicating a role for CLPB3 in disentangling protein aggregates from there.

## Introduction

The casein lytic proteinase/heat shock protein 100 (Clp/Hsp100) chaperones belong to the large family of AAA+ proteins (ATPases associated with various cellular activities) (Neuwald *et al*., 1999). Clp/Hsp100 proteins are divided into two classes, with class II members containing one (ClpX, ClpY), and class I members two AAA+ modules in tandem (ClpA to E) (Schirmer *et al*., 1996). The AAA+ module features the Walker A and B motifs. Clp/Hsp100 assemble into homohexameric rings with a central pore through which protein substrates are threaded (Guo *et al*., 2002; Lee *et al*., 2003; Mogk *et al*., 2018; Mogk *et al*., 2015). ClpA, ClpC, ClpE and ClpX contain a conserved tripeptide [LIV]-G-[FL] crucial for binding to an associated protease like ClpP that is lacking in ClpB/Hsp101 and ClpY family members (Kim *et al*., 2001). ClpB/Hsp101 family members contain two additional domains: the N-terminal domain and the middle domain, which forms a coiled-coil structure that is inserted in the first AAA+ module (Mogk *et al*., 2015). The threading activity of *E. coli* ClpB can initiate at the N-or C-termini or at internal sites of substrate proteins in protein aggregates such that entire peptide loops are translocated through the pore (Avellaneda *et al*., 2020). Translocation is mediated by mobile loops in the central pore that contact the substrate via conserved aromatic residues (Deville *et al*., 2017; Rizo *et al*., 2019). Driven by ATP hydrolysis, the loops move downwards along the translocation channel. Axial staggering of the loops facilitates substrate handover and prevents substrate backsliding. Once displaced to the opposite side of the ClpB hexameric ring, the substrate can fold to the native state by itself or aided by Hsp70 and/or chaperonins (Mogk *et al*., 2018).

The ability of cytosolic ClpB/Hsp100 members to disentangle individual proteins from aggregates has been shown to be crucial for the resolving of protein aggregates formed upon heat stress in yeast and in Arabidopsis (Agarwal *et al*., 2003; McLoughlin *et al*., 2016; Parsell *et al*., 1994). Eliminating cytosolic HSP101 activity in land plants had no effect on plant growth under ambient temperatures but resulted in reduced basal and acquired thermotolerance and, *vice versa*, increasing HSP101 levels resulted in enhanced basal and acquired thermotolerance (Hong and Vierling, 2000, 2001; Katiyar-Agarwal *et al*., 2003; Nieto-Sotelo *et al*., 2002; Queitsch *et al*., 2000). Deletion of the single copy-gene encoding ClpB in *Synechococcus sp*. had no phenotype under optimal growth conditions, either, and did not affect basal thermotolerance but strongly impaired the capacity of the mutant to develop thermotolerance (Eriksson and Clarke, 1996, 2000). Hence, to survive heat stress it is crucial that cells resolve heat-induced cytosolic protein aggregates during recovery from heat. The engineering of yeast cytosolic Hsp104 to deliver substrates directly to an associated peptidase abolished thermotolerance, suggesting that for thermotolerance a reactivation of aggregated cytosolic proteins is required, not just their removal (Weibezahn *et al*., 2004).

In plants, chloroplast ClpB (termed CLPB3, ClpB-p, or APG6) and mitochondrial ClpB (termed CLPB4 or ClpB-m) are both derived from cyanobacterial ClpB (Lee *et al*., 2007; Mishra and Grover, 2016). Suppressing the expression of chloroplast CLPB3 in tomato did not result in visible phenotypes under optimal growth conditions but strongly impaired acquired thermotolerance. This suggests that also in the chloroplast protein aggregates, formed under heat stress, must be resolved in the recovery phase to promote survival (Yang *et al*., 2006). Interestingly, Arabidopsis *clpb3* knock-out mutants under ambient conditions were pale green with smaller and rounder chloroplasts lacking starch grains and showing an abnormal development of thylakoid membranes when compared with the wild type (Lee *et al*., 2007; Myouga *et al*., 2006; Zybailov *et al*., 2009). Accordingly, *clpb3* mutant plants exhibited lower PSII activity and were seedling-lethal if not provided with sucrose, pointing to a role of CLPB3 as a general housekeeping chaperone in chloroplasts, at least in Arabidopsis. Surprisingly, unlike in tomato, thermotolerance was not impaired in Arabidopsis *clpb3* mutants and was not enhanced when CLPB3 was overexpressed (Lee *et al*., 2007; Myouga *et al*., 2006). Nevertheless, recombinantly produced Arabidopsis CLPB3 was found to interact with heat-denatured, aggregated G6PDH and to support the disentangling and refolding of a large part of the protein to the native state in an ATP-dependent reaction *in vitro* (Parcerisa *et al*., 2020). Moreover, Arabidopsis CLPB3 promoted the refolding of aggregation-prone DXS (the rate-determining enzyme for the production of plastidial isoprenoids) under ambient conditions *in vivo* (Llamas *et al*., 2017; Pulido *et al*., 2016).

*Chlamydomonas reinhardtii* has five *CLPB* genes (Schroda and Vallon, 2009). CLPB1, CLPB3, and CLPB4 are expressed at low levels under ambient temperature (CLPB1 lower than CLPB3/4) and accumulate rapidly and with similar kinetics during heat stress at 42°C, with a plateau reached after 2 h at 42°C (Mühlhaus *et al*., 2011). CLPB2 and CLPB5 have not been detected in proteomics studies, lack EST support and it is therefore not sure whether they are produced only under certain conditions or not at all (Schroda and Vallon, 2009). CLPB1 is predicted to be localized to the cytosol, CLPB3 to the chloroplast, while the localization of CLPB4 is not clear. CLPB3, together with HSP22E, HSP22F, HSP22C, VIPP1, VIPP2, and DEG1C, was up-regulated when chloroplasts experienced stresses likely to disturb chloroplast protein homeostasis. These stresses include high light intensities or elevated cellular H_2_O_2_ concentrations (Blaby *et al*., 2015; Nordhues *et al*., 2012; Perlaza *et al*., 2019; Theis *et al*., 2019; Theis *et al*., 2020), the depletion of chloroplast-encoded ClpP (Ramundo *et al*., 2014), the depletion of thylakoid membrane transporters/integrases (Theis *et al*., 2019; Theis *et al*., 2020), the addition of nickel ions (Blaby-Haas *et al*., 2016) or the alkylating agent methyl methanesulfonate (Fauser *et al*., 2022), and the inhibition of chloroplast fatty acid synthesis (Heredia-Martínez *et al*., 2018). These seven chloroplast proteins appear to represent a core set of proteins involved in the coping with disturbed chloroplast protein homeostasis (Perlaza *et al*., 2019; Ramundo *et al*., 2014). Their upregulation appears to be triggered by misfolded/misassembled proteins inducing lipid packing stress in chloroplast membranes that is sensed and coped with by the VIPP1/2 proteins (Kleine *et al*., 2021; Theis *et al*., 2020). HSP22E/F were found to interact with thermolabile stromal proteins and chaperones in heat stressed cells and with VIPP1/2 and stromal HSP70B especially at chloroplast membranes in cells exposed to H_2_O_2_ (Rütgers *et al*., 2017; Theis *et al*., 2020). DEG1C localizes to the stroma and the periphery of thylakoid membranes. Purified DEG1C exhibited high proteolytic activity against unfolded model substrates, which increased with temperature and pH (Theis *et al*., 2019). No functional studies on Chlamydomonas CLPB3 exist so far. Therefore, the aim of this work was to shed light on CLPB3 function, in particular regarding its possible role in maintaining chloroplast protein homeostasis. We show that CLPB3 is crucial for removing protein aggregates in the chloroplast, which contributes to enhanced thermotolerance under conditions of severe heat stress.

## Materials and Methods

### Strains and cultivation conditions

*Chlamydomonas reinhardtii* wild-type strain CC-4533 (*cw15, mt-*) and mutant strains *clpb3-1* (LMJ.RY0402.250132_1) and *clpb3-2* (LMJ.RY0402.104257_1) from the *Chlamydomonas* Library Project (Li *et al*., 2016) were obtained via the *Chlamydomonas* Resource Center (https://www.chlamycollection.org/). Cultures were grown mixotrophically in TAP medium (Kropat *et al*., 2011) on a rotatory shaker at 25°C and ∼40 μmol photons m^-2^ s^-1^. For complementation, *clpb3* mutant cells were transformed via the glass bead method (Kindle, 1990) as described previously (Hammel *et al*., 2020) with the constructs linearized via EcoRV digestion. Transformants were selected on TAP medium containing 100 μg/mL spectinomycin. Cell densities were determined using the Z2 Coulter Counter (Beckman Coulter) or photometrically by optical density measurements at 750 nm (OD_750_). For heat stress experiments, cultures of exponentially growing cells were placed into a water bath heated to 40°C and incubated under agitation and constant illumination at ∼40 μmol photons m^-2^ s^-1^ for 1 h, with subsequent recovery at 25°C for 6 h. For spot tests, cells were grown to a density of 3-5 × 10^6^ cells mL^-1^ and diluted in TAP medium or high salt medium (HSM) such that 10 μl contained 10^4^, 10^3^ or 10^2^ cells. 10 μl of each dilution were spotted onto agar plates containing TAP medium or HSM medium. HSM was prepared according to Sueoka (1960) but using the trace solutions from Kropat *et al*. (2011).

### Extraction of *Chlamydomonas* genomic DNA and verification of the insertion sites

For the extraction of *Chlamydomonas* genomic DNA, 5 mL of exponentially growing cells were pelleted and resuspended in 250 μL water. 250 μL 2× lysis buffer (20 mM Tris-HCl, 40 mM Na_2_EDTA, 1% (w/v) SDS) and 3 μL proteinase K (NEB: P8102S, 20 mg/mL) were added and incubated under agitation at 55°C for 2 h. The lysate was supplemented with 80.9 μL of 5 M NaCl and mixed by vortexing. After the addition of 70 μL pre-warmed CTAB/NaCl (2% (w/v) CTAB; 1.4 M NaCl) lysates were vortexed and incubated under agitation for 10 min at 65°C. Nucleic acids were extracted by addition of 1 volume phenol/chloroform/isoamylalcohol (25:24:1; Roth), mixing the two phases and separating for 5 min at 18,000 g and 4°C. Phenol/chloroform extraction of the aqueous phase was repeated once. An equal volume chloroform/isoamylalcohol (24:1; Roth) was added to the upper phase and the mixture was centrifuged as above. Recovering of the nucleic acid was achieved by precipitating with an equal volume isopropanol. Finally, the pellet was resuspended in TE-buffer (10 mM Tris-HCl, pH 8.0; 1 mM EDTA). 1 ng was used for PCR. Validation of the *aphVIII* cassette insertion site within the genes of the CLiP mutant lines was performed using the specific primers listed in Table S1 according to the manual provided by the CLiP (Li *et al*., 2016). Amplified products were analysed by agarose gel electrophoresis. Electrophoresed DNA was stained with Gelred (Biotium) or HDGreen Plus DNA Stain (INTAS Science Imaging) and visualized under UV light using a gel documentation system (FUSION-FX7 Advance™ imaging system (PeqLab) / ECL ChemoStar V90D+ (INTAS Science Imaging).

### Cloning, production, and purification of recombinant CLPB3

The CLPB3 coding region lacking the chloroplast transit peptide was amplified by PCR from EST clone AV631848 (Asamizu *et al*., 2000) with primer CLPB3-Eco and CLPB3-Hind. The 2827-bp PCR product was digested with EcoRI and HindIII and cloned into EcoRI-HindIII-digested pETDuet-1 vector (Novagen) lacking two nucleotides upstream from the BamHI site, producing pMS976. CLPB3 was expressed with an N-terminal hexahistidine tag in *E. coli* Rosetta cells after induction with 1 mM IPTG for 16 h at 20°C and purified by cobalt-nitrilotriacetic acid affinity chromatography according to the manufacturer’s instructions (G-Biosciences), including a washing step with 5 mM ATP. Eluted CLPB3 was gel filtrated using an Enrich SEC650 column. Fractions containing CLPB3 were pooled and concentrated in Amicon® Ultra-4 Centrifugal Filter Units (Ultracel®-3K, Merck Millipore Ltd), with a subsequent buffer exchange to 6 M Urea, 50 mM NaCl, 20 mM Tris-HCl, pH 7.5. Protein amounts were determined via a NanoDrop 2000 (ThermoFischer Scientific) Proteins were frozen in liquid nitrogen and stored at -80°C. 2.6 mg of the protein was used for the immunization of a rabbit via the 3-month standard immunization protocol of Bioscience bj-diagnostik (Göttingen).

### Plasmid constructs for the complementation of *clpb3* mutants

The genomic *CLPB3* gene, ranging from start to stop codon and including all introns except for introns 7, 8 and 11, was synthesised in three fragments with flanking BsaI restriction sites. The fragments were cloned into the pTwist Kan High Copy vector by Twist Bioscience, resulting in three level 0 constructs L0-*CLPB3*-up containing a 1933-bp fragment (pMBS495), L0-*CLPB3*-down1 with a 1713-bp fragment (pMBS496), and L0-*CLPB3*-down2 with a 1010-bp fragment (pMBS497). All three level 0 constructs were combined with plasmids pCM0-020 (*HSP70A/RBCS2* promoter + 5’UTR), pCM0-100 (3xHA), and pCM0-119 (*RPL23* 3’UTR) from the *Chlamydomonas* MoClo kit (Crozet *et al*., 2018) as well as with destination vector pICH47742 (Weber *et al*., 2011), digested with BsaI and ligated to generate level 1 construct pMBS587 harbouring the full *CLPB3* transcription unit encoding a C-terminal 3xHA-tag. The level 1 construct was then combined with pCM1-01 (level 1 construct with the *aadA* gene conferring resistance to spectinomycin flanked by the *PSAD* promoter and terminator (Crozet *et al*., 2018)), with plasmid pICH41744 containing the proper end-linker, and with destination vector pAGM4673 (Weber *et al*., 2011), digested with BbsI, and ligated to yield level 2 construct pMBS588. Correct cloning was verified by Sanger sequencing.

### Protein analyses

Protein extractions, SDS-PAGE, semi-dry blotting and immunodetections were carried out as described previously (Liu *et al*., 2005; Schulz-Raffelt *et al*., 2007). Sample amounts loaded were based on protein (Bradford, 1976) or chlorophyll concentrations (Porra *et al*., 1989). Immunodetection was performed using enhanced chemiluminescence (ECL) and the FUSION-FX7 Advance™imaging system (PEQLAB) or ECL ChemoStar V90D+ (INTAS Science Imaging). Antisera used were against CLPB3 (this study), CGE1 (Schroda *et al*., 2001), HSP22E/F (Rütgers *et al*., 2017), DEG1C (Theis *et al*., 2019), TIG1 and PRPL1 (Ries *et al*., 2017), CPN60A (Westrich *et al*., 2021), and the HA-tag (Sigma-Aldrich H3663). Anti-rabbit-HRP (Sigma-Aldrich) and anti-mouse-HRP (Santa Cruz Biotechnology sc-2031) were used as secondary antibodies. Densitometric band quantifications after immunodetections were done by the FUSIONCapt Advance program (PEQLAB).

### Isolation of protein aggregates

Protein aggregates were isolated as described previously (Koplin *et al*., 2010) with minor modifications. Briefly, *Chlamydomonas* cells were grown to a density of approximately 5 × 10^6^ cells/ml and a total of 2 × 10^8^ cells were used. Cells were supplemented with sodium azide at a final concentration of 0.002% and harvested by centrifugation at 3,500 g for 2 min at 4°C, and cell pellets were frozen in liquid nitrogen and stored at −80°C. Cell pellets were thawed on ice and resuspended in lysis buffer (20 mM sodium phosphate pH 6.8, 10 mM DTT, 1 mM EDTA, and 0.25× protease inhibitor cocktail [Roche]). Cells were sonicated on ice and centrifuged for 10 min at 500 g and 4 °C to remove intact cells and cell debris. Protein concentrations in the supernatant were measured by the Bradford assay (Bradford, 1976), and samples were diluted to match the sample with the lowest protein concentration. Samples for total input were taken and supplemented with 2× Laemmli sample buffer (125 mM Tris–HCl pH 6.8, 20% glycerol, 4% SDS, 0.1 M DTT, and 0.005% bromophenol blue). Samples were then centrifuged for 30 min at 19,000 g and 4°C. Pellets were washed 4 times by repeated sonication with washing buffer containing 20 mM sodium phosphate pH 6.8 and 2% Nonidet-P40 and centrifugation for 30 min at 19,000 g and 4°C. At last, pellets were dissolved in 1× Laemmli sample buffer containing 3 M urea. Samples were separated on 12% SDS-polyacrylamide gels followed by Coomassie staining or immunoblotting.

### Blue native PAGE analysis

Blue native (BN) PAGE with whole cell proteins was carried out according to published protocols (Schagger *et al*., 1994; Schagger and von Jagow, 1991) with minor modifications. Briefly, cells were exposed for 1 h to 41°C heat shock with a subsequent 6 h recovery at 25°C as described above. Approximately 10^8^ cells were harvested by centrifugation, washed with TMK buffer (10 mM Tris-HCl, pH 6.8, 10 mM MgCl_2_, 20 mM KCl), and resuspended in 500 μL ACA buffer (750 mM ε-aminocaproic acid, 50 mM Bis-Tris pH 7.0 and 0.5 mM EDTA) supplemented with 0.25× protease inhibitor (Roche). Cells were broken by sonication. Intact cells and cell debris were removed by centrifugation for 5 min at 300 g and 4°C. Whole cell lysates (equivalent to 0.25 μg/μL of chlorophyll) were solubilized for 20 min with 1% (w/v) β-dodecyl maltoside (Roth) on ice in the dark and insolubilized material was precipitated by centrifugation at 18500 g for 10 min at 4°C. Afterwards, supernatants were supplemented with native sample buffer (750 mM ε-aminocaproic acid and 5% (w/v) Coomassie Brilliant Blue G250) and separated on a 4-15% (w/v) blue-native polyacrylamide gel followed by immunoblotting.

### Microscopy

For immunofluorescence microscopy, cells were fixed and stained as described previously (Uniacke *et al*., 2011) with minor modifications: microscopy slides were washed three times with 100% ethanol and coated with 0.1% poly-L-lysine. Cells were fixed with 4% formaldehyde for at least 1 h at 4°C on an overhead rotator. Aliquots of 40 μL cell suspension were allowed to adhere to the microscope slides for 15 min at 25°C, followed by incubation in 100% methanol for 6 min at -20°C. Afterwards, slides were washed five times with phosphate-buffered saline (PBS). Cells were permeabilized by incubating the slides with 2% Triton X-100 in PBS for 10 min at 25°C. Slides were washed three times with PBS containing 5 mM MgCl_2_ and with PBS-BSA (PBS, 1% BSA) for at least 30 min at 25°C. Slides were incubated over night at 4°C with antisera against HSP22EF and the HA-tag in 1:1000 dilutions in PBS-BSA. Slides were then washed five times with PBS-BSA at 25°C followed by incubation in a 1:200 dilution of the tetramethylrhodamine-isothiocyanate-labelled goat anti-rabbit antibody (TRITC, Sigma-Aldrich) and fluorescein isothiocyanate-labelled goat anti-mouse antibody (FITC, Sigma-Aldrich) in PBS-BSA for 1.5 h at 25°C in the dark. Finally, the slides were washed five times with PBS and mounting solution containing 4’,6-diamidino-2-phenylindole (DAPI; Vectashield; Vector Laboratories) was dispersed over the cells. HSP22EF and the HA-tag images were captured with a Zeiss LSM880 AxioObserver confocal laser scanning microscope equipped with a Zeiss C-Apochromat 40x/1.2 W AutoCorr M27 water-immersion objective. Fluorescent signals of FITC (excitation/emission 488 nm/491–589 nm) and TRITC (excitation/emission 633 nm/647–721 nm) were processed using the Zeiss software ZEN 2.3 or ImageJ. Light microscopy images were taken with an Olympus BX53 microscope.

### Chlorophyll fluorescence measurements

Chlorophyll fluorescence was measured using a pulse amplitude-modulated Mini-PAM fluorometer (Mini-PAM, H. Walz, Effeltrich, Germany) essentially according to the manufacturer’s protocol after 3 min of dark adaptation (1 s saturating pulse of 6000 μmol photons m-2 s-1, gain = 4).

## Results

The Chlamydomonas *CLPB3* gene encodes a preprotein with 1043 amino acids of which the N-terminal 115 ones are predicted to serve as a chloroplast transit peptide (Fig. S1). The mature CLPB3 protein has a mass of 101 kDa and shares 54% identical and 72% similar residues with *E. coli* ClpB and 68% identical and 82% similar residues with mature *Arabidopsis* ClpB3. We produced mature *Chlamydomonas* CLPB3 recombinantly in *E. coli* with an N-terminal hexa-histidine tag (Figs. S1 and S2) and raised a polyclonal antibody. The antibody revealed that CLPB3 is expressed constitutively in *Chlamydomonas* as a protein with an apparent molecular mass of ∼102 kDa that migrated little below the full-length recombinant protein, indicating processing of the transit peptide at the predicted site (Fig. 1; Fig. S1). We observed some degradation of the recombinant protein. Quantification of the immunoblot signals (including the degradation products) revealed that CLPB3 accounts for about 0.2% ± 0.024% (SD, n = 3) of total cell proteins.

**Fig. 1.**
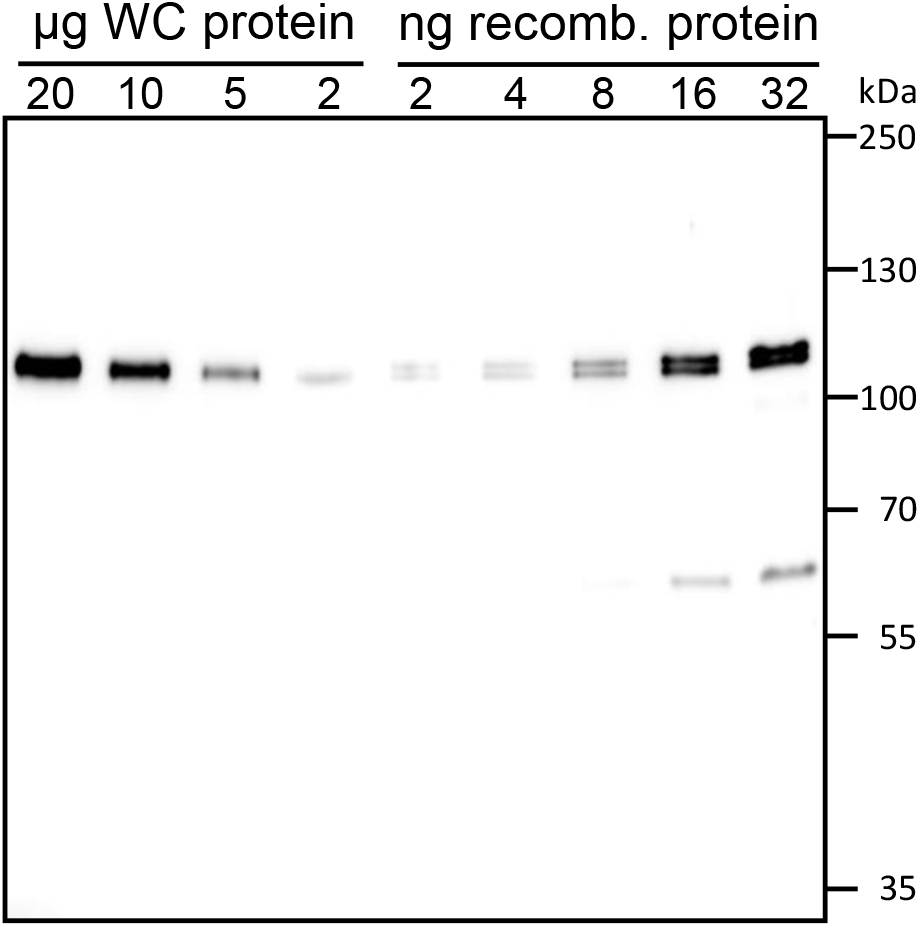
Quantification of CLPB3. The indicated amounts of whole-cell (WC) protein from *Chlamydomonas* wild type grown at 25°C and of CLPB3 produced recombinantly in *E. coli* were separated on a 8% SDS-polyacrylamide gel and analyzed by immunoblotting using an antibody raised against *Chlamydomonas* CLPB3.

### Two *clpb3* mutants accumulate a truncated form of CLPB3 and less CLPB3

To obtain insights into the function of CLPB3 in *Chlamydomonas*, we ordered two *clpb3* mutants from the Chlamydomonas library project (CLiP) (Li *et al*., 2016) with insertions of the mutagenesis cassette in exon 12 (*clpb3-1*) and in intron 4 (*clpb3-2*) (Fig. 2a). We could amplify both flanking regions of the cassette in the *clpb3-2* mutant and the flanking region 5’ of the cassette in the *clpb3-1* mutant (Fig. S3). However, we could not amplify the flanking region 3’ of the cassette in the *clpb3-1* mutant, even with staggered flanking primers, but we could show that the cassette is intact. Most likely, additional DNA sequences were inserted between the 3’ end of the cassette and the insertion site in the *CLPB3* gene, which is not uncommon for *Chlamydomonas* insertional mutants (Spaniol *et al*., 2022; Zhang *et al*., 2014).

**Fig. 2.**
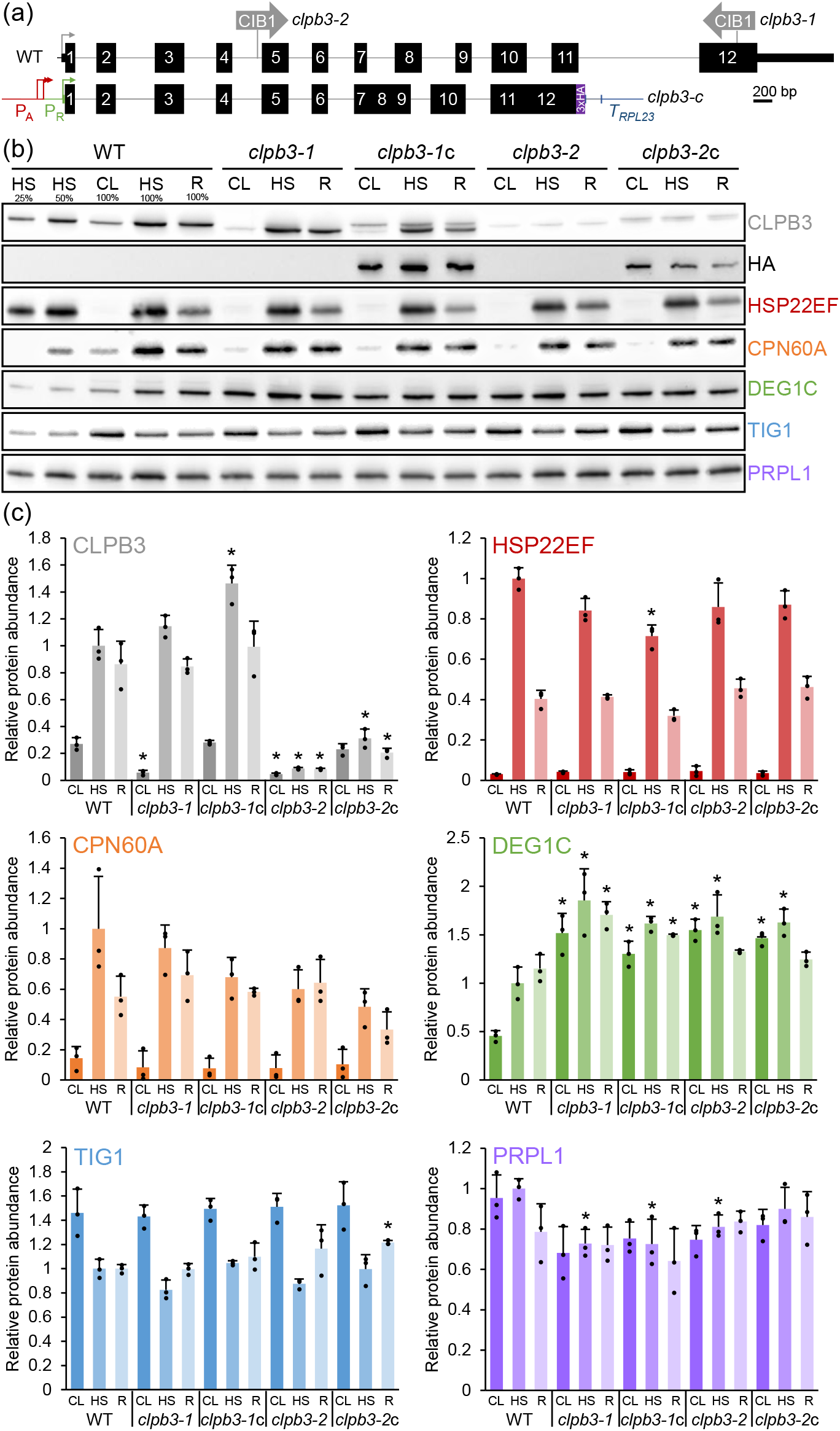
Accumulation of chloroplast proteins with roles in protein homeostasis in wild type, *clpb3* mutants and complemented lines. (a) Structure of the *Chlamydomonas CLPB3* gene, insertion sites of the CIB1 cassette in the *clpb3-1* and *clpb3-2* mutants, and construct for complementation. Protein coding regions are drawn as black boxes, untranslated regions as bars, and introns, promoters and intergenic regions as thin lines. Arrows indicate transcriptional start sites. WT – wild type; *clpb3-*c *–* complemented mutants; P_A_, P_R_ – *HSP70A* and *RBCS2* promoters; T_RPL23_ – *RPL23* terminator. (b) Immunoblot analysis of the accumulation of CLPB3 and selected chloroplast proteins. Cells were grown in continuous light at 25°C (CL), exposed to 41°C for 1 h (HS), and allowed to recover at 25°C for 6 h after the heat treatment (R). 10 μg of whole-cell proteins (100%) were analyzed. (c) Quantification of immunoblot analyses. Values are means from three independent experiments (including two technical ones for CLPB3 and HSP22EF), normalized first by the median of all signals obtained with a particular antibody in the same experiment, and then by the mean signal of the heat-stressed wild type. Error bars represent standard deviation. Asterisks indicate significant differences with respect to the WT (two-tailed, unpaired t-test with Bonferroni-Holm correction, P < 0.05). The absence of an asterisk means that there were no significant differences.

We first analyzed CLPB3 levels in the mutants and in wild type under ambient conditions, after a 60-min exposure to 40°C, and after 6 h of recovery from heat stress. As shown in Figs.2b and 2c, CLPB3 levels in the wild type increased 4-fold during the heat treatment and declined by ∼14% during recovery, corroborating findings from a large-scale proteomics study (Mühlhaus *et al*., 2011). We found two putative heat shock elements (HSEs) about 60 nt and 90 nt upstream of a putative TATA box in the *CLPB3* gene that show a degree of degeneration typical for HSEs in Chlamydomonas (Fig. S4) (Lodha *et al*., 2008). These HSEs most likely are driving the heat-induced expression of the *CLPB3* gene via heat shock transcription factor HSF1 (Schulz-Raffelt *et al*., 2007).

In both mutants, CLPB3 accumulated only to ∼20% of wild-type levels under ambient conditions. While in the *clpb3-1* mutant CLPB3 after heat treatment and recovery accumulated like in wild type, CLPB3 levels barely increased in the *clpb3-2* mutant. Apparently, intron splicing in this mutant is impaired and results in overall lower protein production. CLPB3 in the *clpb3-1* mutant migrated with a slightly smaller apparent molecular mass of ∼96 kDa than in the wild type and the *clpb3-2* mutant, in line with the predicted truncation of its C-terminus (Fig. S1). Unlike in Arabidopsis *clpb3* mutants, we observed no obvious phenotypes in chloroplast development or photosystem II activity in the two Chlamydomonas *clpb3* mutants (Supplemental Fig. S5).

### CLPB3 abundance in the mutants can be partially restored in complemented lines

To complement the mutants, we synthesized the genomic sequence encoding the entire CLPB3 protein as a level 0 part for the Modular Cloning system (Crozet *et al*., 2018). All introns were kept, except introns 7, 8, and 11 because they contain highly repetitive sequences. The *CLPB3* genomic sequence was then assembled into a level 1 module with the *HSP70A-RBCS2* promoter, *RPL23* terminator, and a sequence encoding a C-terminal 3xHA tag (Fig. 2a). We used the constitutive *HSP70A-RBCS2* promoter because it strongly enhances chances for transgene expression in Chlamydomonas (Strenkert *et al*., 2013). Since cytosolic HSP101 fused C-terminally to GFP or to a Strep tag was fully functional (McLoughlin *et al*., 2016; McLoughlin *et al*., 2019), we did not expect the C-terminal 3xHA sequence to interfere with CLPB3 function. After adding a spectinomycin resistance cassette in a level 2 device, the latter was transformed into both mutants and spectinomycin-resistant transformants were screened using antibodies against CLPB3 and the HA epitope (Fig. S6). Despite using the *HSP70A-RBCS2* promoter, less than 10% of the transformants expressed HA-tagged CLPB3 to clearly detectable levels. Under ambient conditions, the best-expressing transformants accumulated CLPB3 to wild-type levels (*clpb3-1*c) or to 85% of wild type levels (*clpb3-2*c) (Fig. 2b and 2c). After heat shock, CLPB3 levels in *clpb3-1*c exceeded those in the wild type by ∼1.5-fold, while CLPB3 levels in *clpb3-2*c amounted to only ∼30% of wild-type levels.

### Loss of function of CLPB3 results in strongly elevated DEG1C levels

We next analyzed the accumulation of selected proteins involved in chloroplast protein homeostasis (protein biosynthesis, folding, and degradation) to get an idea whether their accumulation was affected by the reduced CLPB3 levels in the mutants. Chloroplast chaperones CPN60A and HSP22E/F (Bai *et al*., 2015; Rütgers *et al*., 2017) strongly accumulated after heat stress and declined after recovery and thus behaved similar to CLPB3, with little differences between mutants and wild type (Fig. 2b and 2c). Levels of trigger factor TIG1, a thermolabile chaperone involved in protein biogenesis (Ries *et al*., 2017; Rohr *et al*., 2019; Rütgers *et al*., 2017), declined by ∼30% after heat stress in the wild type. There was a trend of a more pronounced decrease in both mutants that appeared to be relieved in the complemented lines. Similarly, levels of chloroplast ribosome subunit PRPL1 (Ries *et al*., 2017) appeared to be overall lower in the mutants as compared to the wild type, with some restoration of PRPL1 levels, especially in the *clpb3-2*c line. The most striking difference between *clpb3* mutants and wild type was observed for stromal protease DEG1C (Theis *et al*., 2019). DEG1C accumulated to much higher levels in the mutants compared to the wild type under all conditions (more than 3-fold under ambient conditions), with a trend towards restoration of lower DEG1C levels, especially in the *clpb3-1*c line (Fig. 2b and 2c). To substantiate these findings, we dedicatedly investigated the accumulation of DEG1C in wild type, *clpb3-2* mutant and *clbb3-2*c only under ambient conditions (Fig. 3). Here the *clpb3-2* mutant accumulated 2.3-fold higher DEG1C levels than the wild type, which were reduced to 1.5-fold higher levels in the complemented line *clbb3-2*c. The lack of full complementation can be explained by the accumulation of CLPB3 to only ∼80% of wild-type levels.

**Fig. 3.**
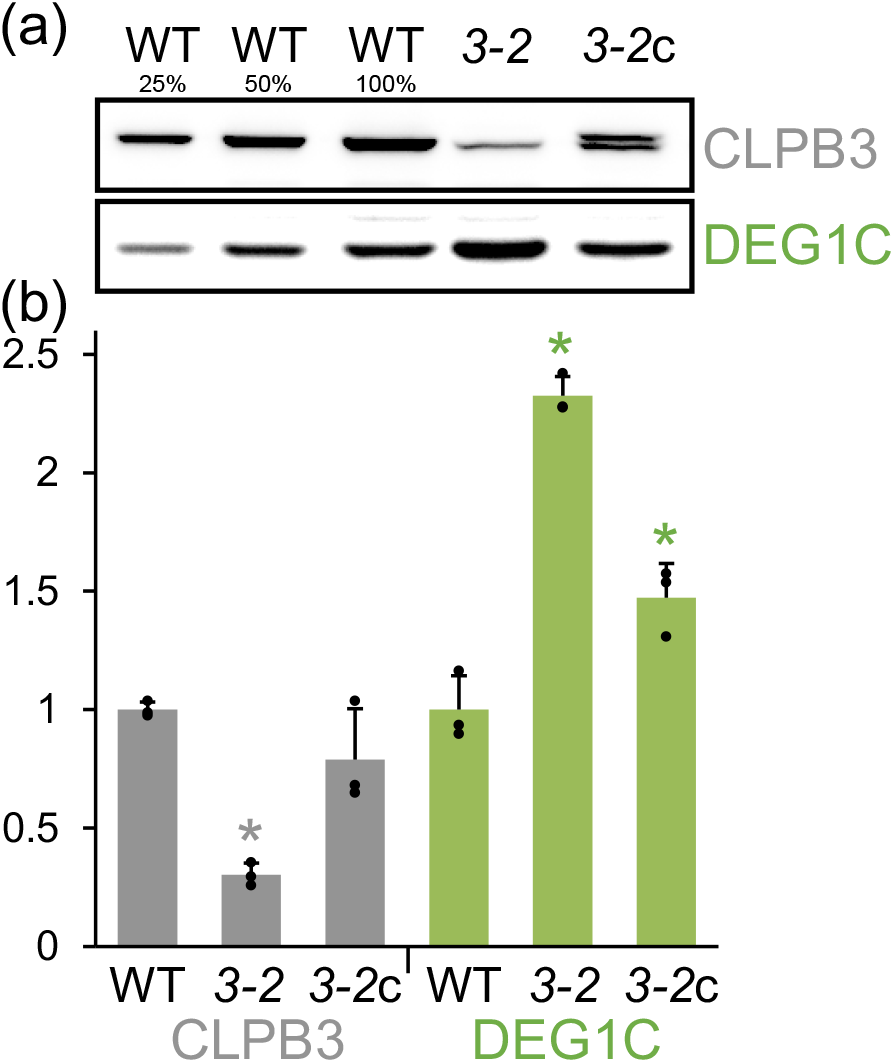
Restoration of DEG1C accumulation in complemented mutant line *clpb3-2*c. (a) Immunoblot analysis of CLPB3 and DEG1C accumulation in wild type (WT), *clpb3-2* mutant (*3-2*) and complemented mutant *clpb3-2*c (*3-2*c). Cells were grown in continuous light at 25°C. 10 μg of whole-cell proteins (100%) were analyzed. (b) Quantification of immunoblot analyses as described for Fig. 1c with normalization on protein levels in WT. Error bars represent standard deviation, n = 3. Asterisks indicate significant differences with respect to the WT (two-tailed, unpaired t-test with Bonferroni-Holm correction, P < 0.05). The absence of an asterisk means that there were no significant differences.

### CLPB3 partitions into aggregates of high molecular weight after heat stress

To assess the oligomeric state of CLPB3, we subjected wild type, *clpb3* mutants and complemented lines to the same heat shock/recovery regime as before and analyzed whole-cell proteins by BN-PAGE and immunoblotting. We detected specific signals for CLPB3 that we assigned to monomers and aggregates of high molecular weight (Fig. 4). Although a signal was observed at the height of photosystem (PS) I that could correspond to CLPB3 hexamers (PS I has a molecular mass of ∼600 kDa (Amunts *et al*., 2007)), the equal intensity of this signal in all lines rather argues for a cross-reaction of the CLPB3 antibody with a PSI subunit. In wild type, the signals for monomers and aggregates increased strongly after heat stress and remained strong after the recovery phase. In the *clpb3-1* mutant the monomer was virtually absent under all conditions, while a very strong signal was detected in aggregates after heat shock and recovery. The same pattern was observed also in the *clpb3-1*c line, but there the transgenic CLPB3 monomer was detected. Low levels of the monomer were detected in both, *clpb3-2* mutant and complemented line *clpb3-2*c. Both lines exhibited much weaker signals for CLPB3 in aggregates than observed in wild type, *clpb3-1*, and *clpb3-1*c.

**Fig. 4.**
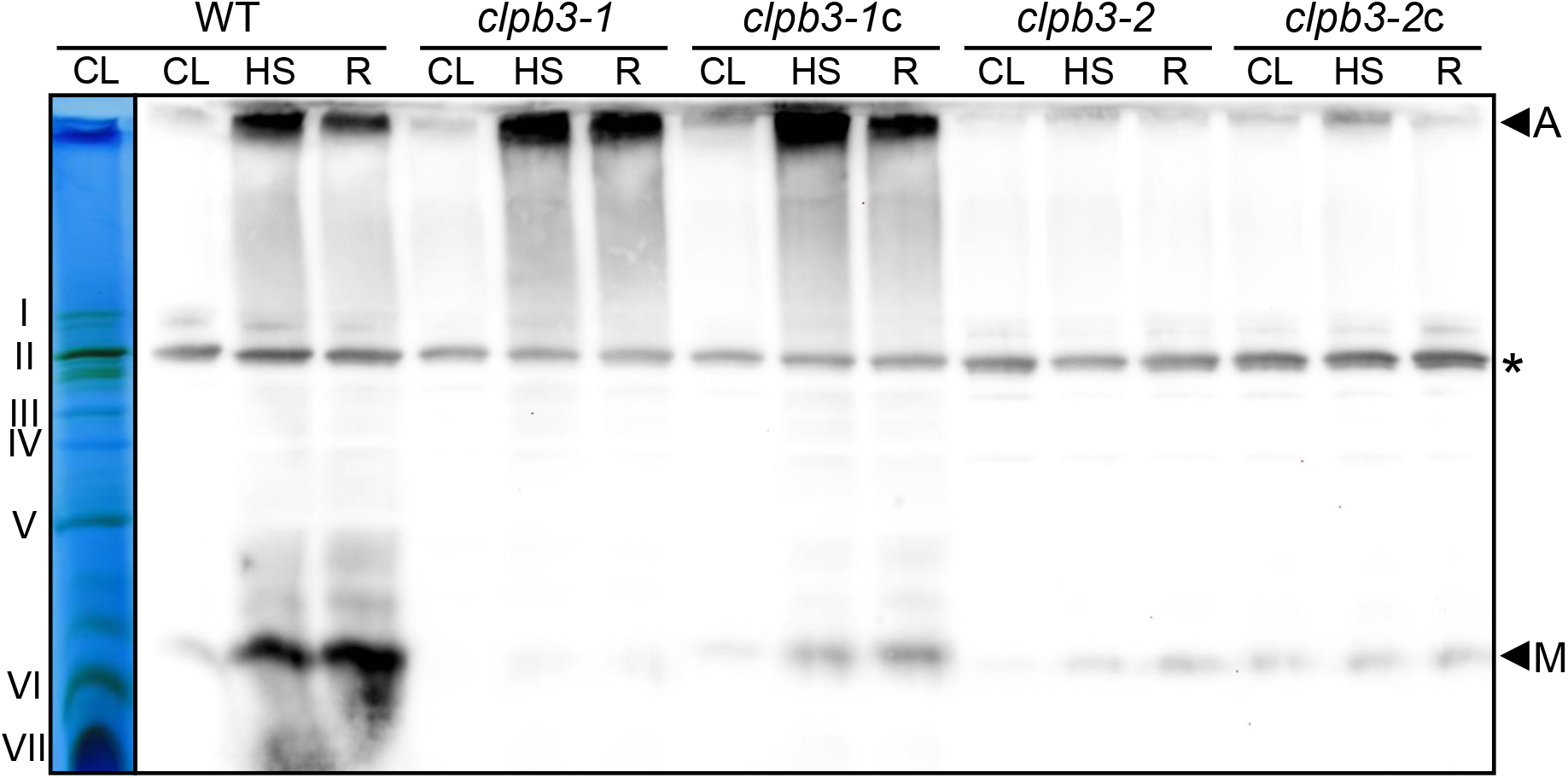
Analysis of the oligomeric state of CLPB3. Whole-cell proteins from wild-type (WT), *clpb3* mutants and complemented lines exposed to the heat shock/recovery regime used in Fig. 2b were solubilized with 1% β-DDM and subjected to BN-PAGE. A lane of the gel after the run is shown at the left with PSII supercomplexes (I+II), PSI-LHCI (II), PSII dimers (III), ATP synthase (IV), PSII monomers/Cyt b6*f* complex (V), LHCII trimers (VI) and LHCII monomers (VII) visible as prominent bands. On the right is an immunoblot of the gel decorated with antibodies against CLPB3. A – Aggregates; M – CLPB3 monomers. The asterisk indicates a protein, presumably of PSI, that cross-reacts with the CLPB3 antibody.

### CLPB3 and HSP22EF localize in stromal foci and to the area occupied by the thylakoid membrane system, respectively

We next employed immunofluorescence to localize CLPB3 in cells of complemented lines exposed to the same heat shock/recovery regime as used before (*clpb3-2*c) or to a 60-min heat shock only (*clpb3-1*c). Since we expected a colocalization of HSP22E/F and CLPB3 in aggregates, we employed mouse antibodies against the HA tag to detect transgenic CLPB3 and rabbit antibodies against HSP22E/F on the same cells. In all cells, HSP22E/F was weakly detectable under ambient conditions and gave rise to strong signals in the chloroplast after heat shock and recovery (Fig. 5), corroborating earlier findings (Rütgers *et al*., 2017). As expected, the HA antibody produced no signals in wild-type cells, but recognized HA-tagged CLPB3 in the complemented lines. In *clpb3-2*c, CLPB3 was evenly dispersed throughout the chloroplast under ambient conditions but partitioned into stromal foci after heat stress. These foci largely vanished after the recovery phase (Fig. 5, top panel). Since transgenic CLPB3-HA levels did not increase during heat stress (Fig. 2), the stromal foci must be formed by the redistribution of existing CLPB3-HA protein under heat and redistribute during recovery. In *clpb3-1*c, CLPB3 localized to stromal foci already under ambient conditions that became stronger and more condensed after heat stress (Fig. 5, bottom panel). To our surprise, in both complemented lines, HSP22E/F and CLPB3 hardly colocalized after heat shock. Rather, CLPB3 stromal foci were more adjacent to HSP22E/F signals that appeared to derive largely from the area occupied by the thylakoid membrane system.

**Fig. 5.**
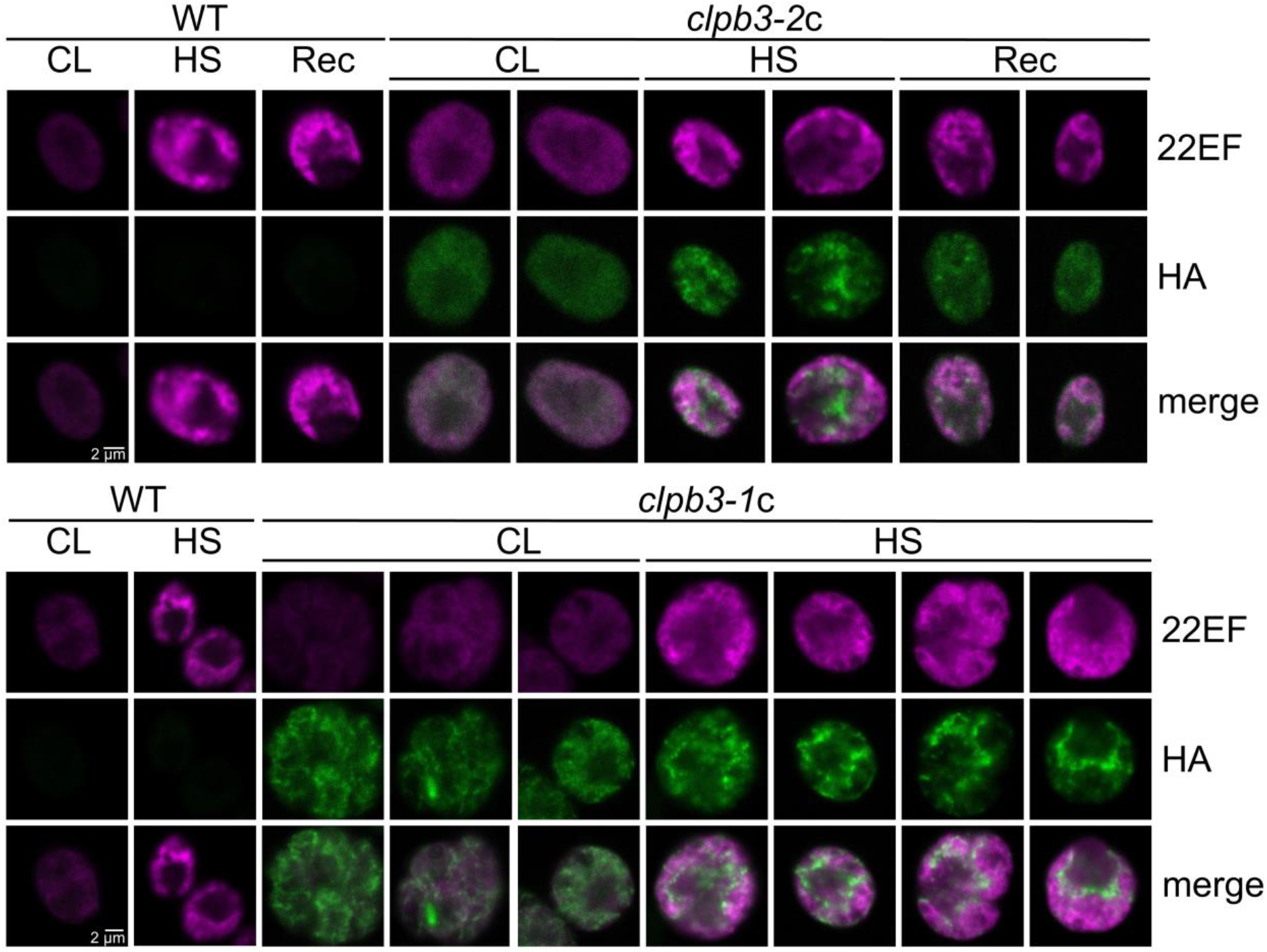
Subcellular localization of CLPB3 and HSP22EF. Cells were exposed to the heat shock/recovery regime used in Fig. 2b (wild type (WT) and *clpb3-2*c) or only to a 60-min heat shock treatment (WT and *clpb3-1*c). HSP22E/F (22EF) and HA-tagged CLPB3 (HA) were detected by immunofluorescence using antibodies against HSP22E/F (magenta) and the HA epitope (green). Merge – overlay of both signals. The scale bar corresponds to 2 μm and applies to all images.

### The removal of aggregated proteins during recovery from heat stress is impaired in *clpb3* mutants

Chloroplast ClpB3 from Arabidopsis was shown to exhibit disaggregase activity *in vitro* (Parcerisa *et al*., 2020). To elucidate a disaggregase function of *Chlamydomonas* CLPB3 *in vivo*, we exposed wild type and *clpb3* mutants to our heat shock/recovery regime and purified protein aggregates, which were analyzed by SDS-PAGE and Coomassie staining (Fig. 6a). In wild type, the abundance of insoluble proteins increased after the heat treatment, but during recovery was reduced to the same levels as before the heat treatment. This was not the case in the *clpb3* mutants, in which insoIuble proteins persisted after recovery. To substantiate this finding, we performed the same experiment with wild type, *clpb3* mutants and complemented lines and analyzed the abundance of CLPB3, HSP22E/F, and thermolabile TIG1 in purified aggregates (Figs. 6b and 6c). In all lines, the three proteins were barely detectable in non-soluble proteins prepared from cells kept under ambient conditions, but accumulated strongly in aggregates collected after the heat treatment. In the wild type, after 6 h recovery from heat, the abundance of CLPB3, HSP22E/F, and TIG1 in aggregates was reduced to 10%, 1.5%, and 3%, respectively, of the levels detected after 60 min heat treatment. In contrast, the *clpb3-2* mutant retained 35%, 61%, and 57%, respectively, of these proteins in aggregates after recovery, and the *clpb3-1* mutant retained as much as 85%, 83%, and 91%, respectively. In the complemented lines, there was a clear trend for an improved removal of the three proteins from aggregates, which in *clpb3-1*c was significant for HSP22E/F and TIG1 (Fig. 6c).

**Fig. 6.**
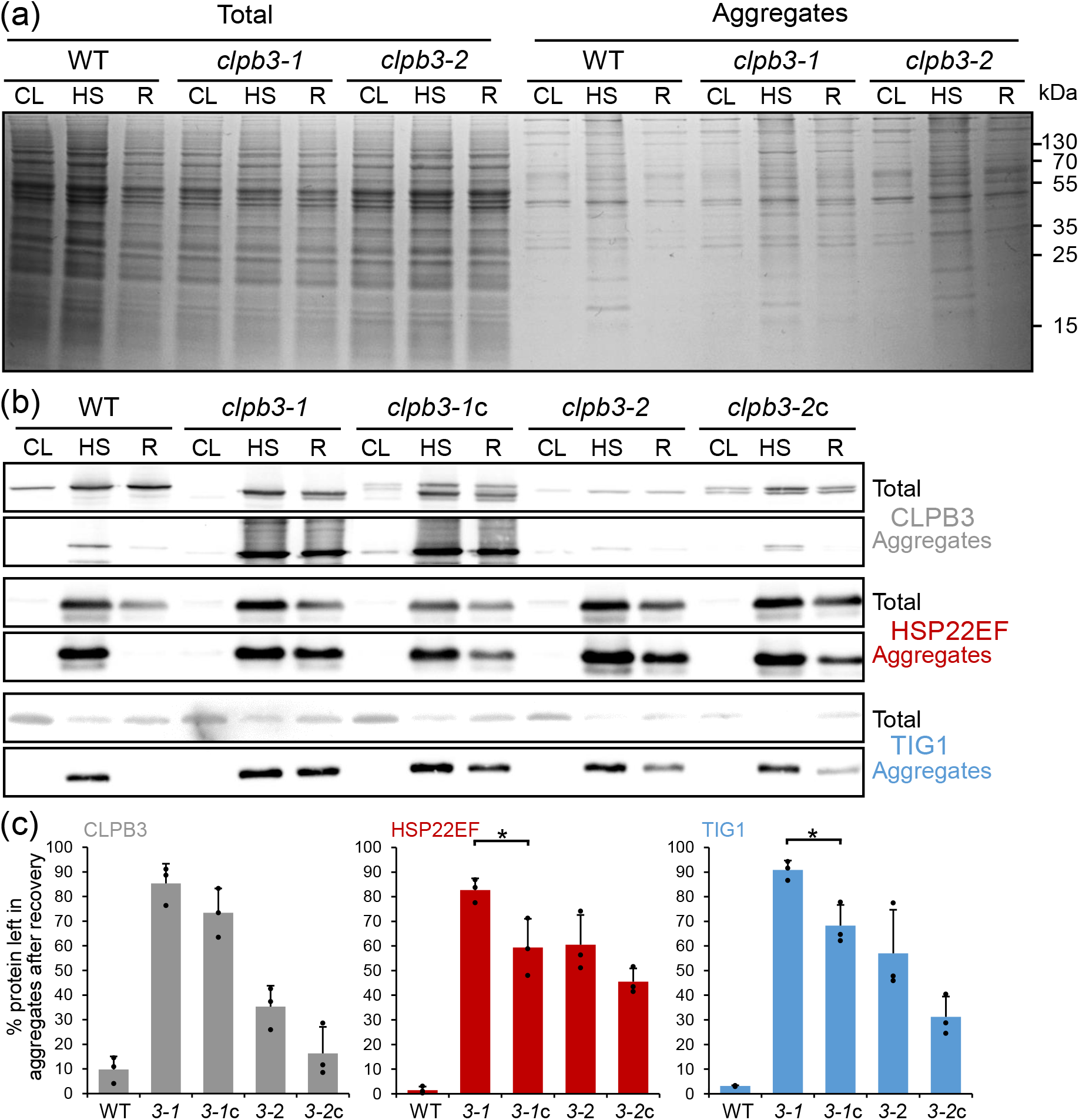
Analysis of aggregate formation and removal in wild type (WT), *clpb3* mutants and complemented lines. (a) Cells were exposed to the heat shock/recovery regime used in Fig. 2b. Total cell proteins and purified aggregates were separated by SDS-PAGE and stained with Coomassie blue. (b) Immunoblot analysis using antibodies against CLPB3, HSP22E/F, and TIG1 on total cell proteins and aggregates. (c) Quantification of the immunoblot analysis shown in (b). Values represent the percentage of protein left in aggregates after 6 h of recovery from three independent experiments. Error bars represent standard deviation. Asterisks indicate significant differences between mutant and its respective complemented line (two-tailed, unpaired t-test, P < 0.05). The absence of an asterisk means that there were no significant differences.

### CLPB3 improves thermotolerance in Chlamydomonas

We wondered whether the impaired ability of the *clpb3* mutants to remove aggregates was associated with a growth phenotype. To test this, we spotted serial dilutions of cultures of wild type, *clpb3* mutants and complemented lines onto agar plates and monitored growth under mixotrophic and photoautotrophic conditions in low and high light, heterotrophic conditions, and repeated prolonged heat stresses under mixotrophic conditions (Fig. 7a). Under all conditions at ambient temperatures, we found no growth phenotype for the *clpb3-2* mutant. The *clpb3-1* mutant exhibited a mild growth phenotype under photoautotrophic conditions in low light and high light, which was ameliorated in the complemented line *clpb3-1*c. Clearly reduced growth was observed for both *clpb3* mutants after repeated prolonged heat stress treatments. This phenotype was ameliorated in *clpb3-2c*, but not in *clpb3-1c*. To substantiate this finding, we exposed cultures of wild type, *clpb3* mutants and complemented lines to 40°C for 72 h and allowed them to recover for 120 h at 25°C. As shown in Fig. 7b, the heat treatment strongly impaired growth of the wild type, but cells resumed growth during the recovery phase. Growth during heat treatment and recovery was abolished in the *clpb3-1* mutant and impaired in the *clpb3-2* mutant when compared with the wild type. This phenotype was ameliorated in both complemented lines, but the wild-type phenotype was not restored. We also determined survival rates for wild-type and the *clpb3-1* mutant after exposure at 41°C for 2 h and found significantly lower survival in the mutant (60%) versus the wild type (89.5%) (Figure 7c). Hence, the growth phenotype after heat stress in the different lines correlated with their ability to remove aggregated proteins during recovery.

**Fig. 7.**
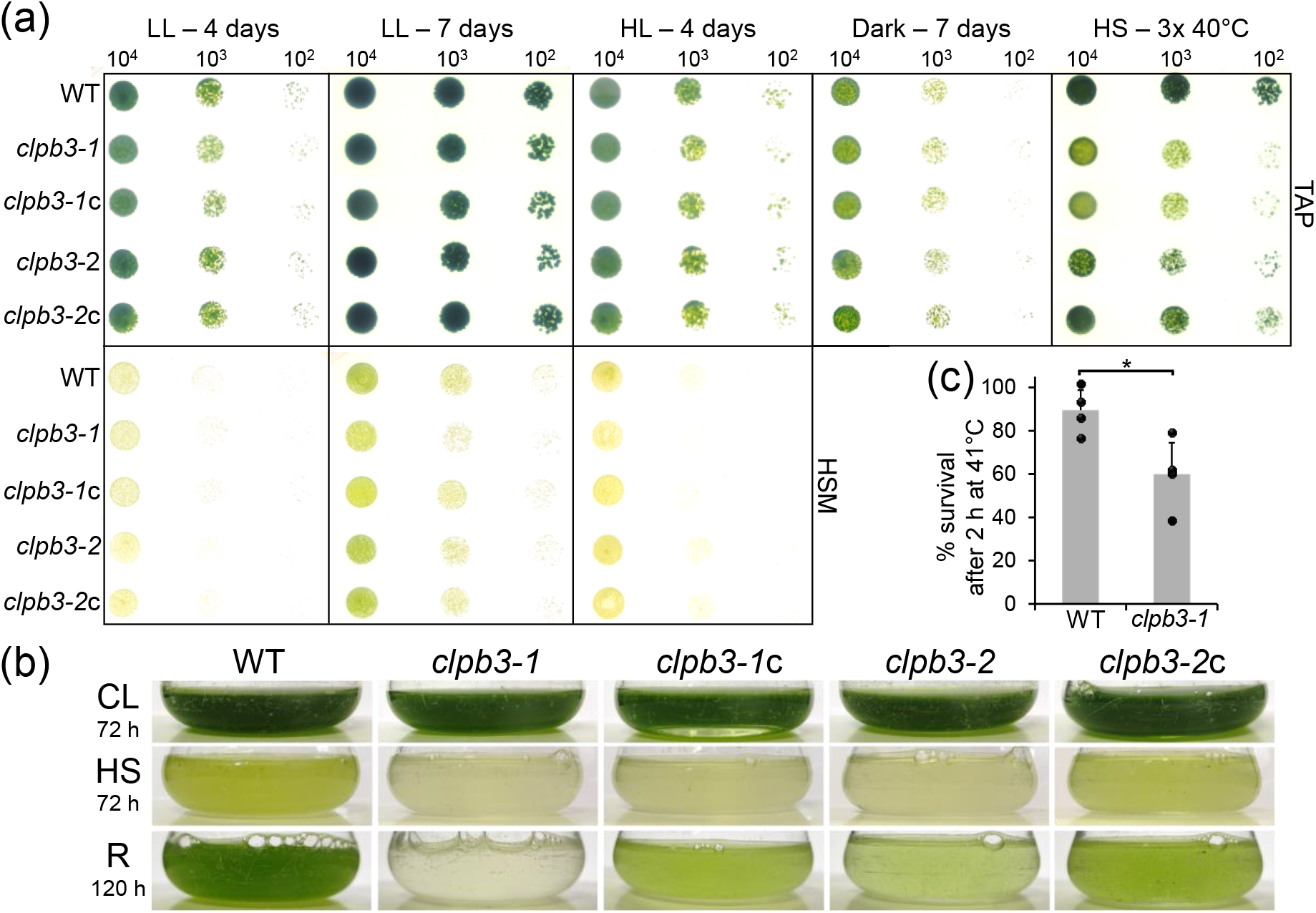
Analysis of growth phenotypes. (a) Wild type (WT), *clpb3* mutants and complemented lines were grown to log phase, diluted, and spotted onto agar plates with the cell numbers indicated. TAP plates were used for monitoring mixotrophic growth (light) or heterotrophic growth (dark), HSM plates for monitoring photoautotrophic growth. LL – low light at 30 μmol photons m^-2^ s^-1^; HL – high light at 600 μmol photons m^-2^ s^-1^; HS – three ∼24 h heat shock exposures at 40°C with ≤24 h recovery in between. (b) Liquid cultures of WT, *clpb3* mutants and complemented lines were grown to log phase, exposed to 40°C for 72 h (HS) and allowed to recover at 25°C for 120 h (R). Before the treatment, part of the culture was diluted and grown at 25°C for 72 h (CL). Shown are pictures of the cultures taken right after the respective treatment. (c) WT and *clpb3-1* mutant were grown to log phase at 25°C and exposed to 41°C for 2 h. Aliquots taken for each condition were diluted, plated on agar plates, and colony-forming units counted after 4 days at 25°C to determine survival rates. Values are from four independent experiments done in triplicates. Error bars represent standard deviation. Differences were significant (two-tailed, unpaired t-test, P < 0.05).

## Discussion

### The resolving of heat-induced protein aggregates by chloroplast CLPB3 is required for thermotolerance in Chlamydomonas

Cytosolic HSP101 is required for the resolving of heat-induced protein aggregates and this activity is essential for basal and acquired thermotolerance in land plants (Agarwal *et al*., 2003; Hong and Vierling, 2000, 2001; Katiyar-Agarwal *et al*., 2003; McLoughlin *et al*., 2016; Nieto-Sotelo *et al*., 2002; Queitsch *et al*., 2000). The situation in chloroplasts is not as clear: while chloroplast CLPB3 is required for thermotolerance in tomato (Yang *et al*., 2006), this is not the case for chloroplast CLPB3 in Arabidopsis (Lee *et al*., 2007; Myouga *et al*., 2006). Nevertheless, Arabidopsis CLPB3 has been shown to be capable of resolving aggregates of model substrate G6PDH *in vitro* and aggregates formed by aggregation-prone DXS *in vivo* (Llamas *et al*., 2017; Parcerisa *et al*., 2020; Pulido *et al*., 2017). Cyanobacterial ClpB, from which chloroplast CLPB3 is derived (Lee *et al*., 2007; Mishra and Grover, 2016), has also been shown to be required for thermotolerance (Eriksson and Clarke, 1996, 2000). In this study we show that chloroplast CLPB3 is required for the resolving of heat-induced protein aggregates in Chlamydomonas (Fig. 6) and that this activity is required for conferring thermotolerance under severe heat stress conditions (Fig. 7). Possibly, a role for chloroplast CLPB3 in conferring thermotolerance in Arabidopsis is obscured by the strong chloroplast development phenotype in Arabidopsis *clpb3* mutants and CLPB3 overexpression lines (Lee *et al*., 2007; Myouga *et al*., 2006; Zybailov *et al*., 2009). An overexpression of compensating chaperones and/or proteases might also play a role.

Chlamydomonas cells appear to compensate the loss of CLPB3 function by upregulating the stromal DEG1C protease and perhaps also by reducing chloroplast protein synthesis capacity, as suggested by a lower abundance of the PRPL1 plastid ribosomal subunit (Figs. 2 and 3). A reduced abundance of cytosolic and chloroplast ribosome subunits was observed in the Chlamydomonas *deg1c* mutant which, however, did not display elevated levels of CLPB3 (Theis *et al*., 2019). The loss of chloroplast CLPB3 function had no effect on the accumulation of other chloroplast chaperones in Chlamydomonas and tomato, including CPN60, HSP70, trigger factor, and sHSPs (Fig. 2) (Yang *et al*., 2006). These observations suggest that a loss of chloroplast disaggregase activity appears to be compensated to some part by lowering the protein synthesis capacity and increasing protease activity rather than by increasing other chaperone systems. However, further research is needed to draw such a conclusion.

### CLPB3 dynamically localizes to stromal foci

We found that heat stress causes CLPB3 to organize in stromal foci by the redistribution of existing protein (Fig. 5). Although HSP22E/F were found in protein aggregates with stromal TIG1 (Fig. 6) and to interact with numerous stromal proteins after heat stress (Rütgers *et al*., 2017), HSP22E/F localized largely to the area occupied by the thylakoid membrane system with little overlap between CLPB3 and HSP22E/F signals (Fig. 5). While the stromal foci formed by CLPB3 in the *clpb3-2*c line largely vanished after the recovery phase, the HSP22E/F signals in the thylakoid membrane area persisted. These results are unexpected, since cytosolic HSP101 and sHSPs in Arabidopsis were found to largely colocalize in cytoplasmic foci (McLoughlin *et al*., 2016; McLoughlin *et al*., 2019). Possibly, HSP22E/F play a dual role during heat stress, with their largest part partitioning to and stabilizing thylakoid membranes and a small part intercalating with stromal proteins in small aggregates for their resolving by CLPB3 in stromal foci. With the bulk HSP22E/F signal coming from the thylakoid system occupied area, this would explain why there is little overlap between the HSP22E/F and CLPB3 signals. Indeed, up to two thirds of Arabidopsis chloroplast Hsp21 have been shown to interact with thylakoid membranes during heat stress (Bernfur *et al*., 2017) and Hsp21 has been shown to stabilize thylakoid membranes and intrinsic protein complexes during heat stress (Chen *et al*., 2017).

In this scenario, the potential functions of HSP22E/F during heat stress would be divided into stabilizing thylakoid membranes and supporting CLPB3-mediated resolution of stromal aggregates. Since the CLPB3 stromal foci look like blobs sitting on HSP22E/F at stroma-exposed regions of the thylakoid system, could CLPB3 play a role there as well? We have previously shown that considerable amounts of HSP22E/F and DEG1C partition to chloroplast membranes upon oxidative stress, where HSP22E/F interact with VIPP1/2 and HSP70B (Theis *et al*., 2020). We proposed that misassembled, unfolded and aggregated proteins might induce lipid packing stress at chloroplast membranes that is sensed by the N-terminal amphipathic α-helix of VIPP2. VIPP2 might then serve as a nucleation point for VIPP1 and HSP22E/F to populate areas suffering from lipid packing stress and prevent membrane leakage. In addition, these proteins might organize membrane domains that serve as interfaces between membrane and soluble chaperones and proteases for the handling of unfolded/aggregated membrane proteins and of aggregates of stromal proteins sticking to the membranes. In this case, CLPB3 might act by resolving such aggregates for refolding or degradation, e.g., via DEG1C. In fact, cytosolic HSP101 has been shown to cooperate with the proteasome system, albeit only on a small subset of aggregated proteins, while refolding was the preferred path (McLoughlin *et al*., 2019). Perhaps membrane proteins threaded through the CLPB3 pore might even be handed over to the ALB3 integrase for reinsertion into the membrane to favor the refolding of membrane proteins over their degradation? Definitely, more work is required to provide evidence for such a bold hypothesis. It is nevertheless attractive, as it provides a coherent function for the main players of the “chloroplast unfolded protein response” regulon, VIPP1/2, HSP22E/F, DEG1C, and CLPB3.

### Despite being abundant under ambient conditions, Chlamydomonas CLPB3 appears not to be required for chloroplast development

We estimated chloroplast CLPB3 to account for ∼0.2% of total cell proteins (Fig. 1). In comparison, the only Hsp70 chaperone in the Chlamydomonas chloroplast, HSP70B, makes up ∼0.19% of total cell proteins (Liu *et al*., 2007). When considering the molar masses, this results in a ratio of 1.4 HSP70B monomers per CLPB3 monomer or about ten HSP70B monomers per CLPB3 hexamer. Upon heat stress, the abundance of CLPB3 increases ∼4-fold, while that of HSP70B increases ∼2.5-fold (Fig. 2) (Mühlhaus *et al*., 2011). Hence, Chlamydomonas CLPB3 is a rather abundant chloroplast protein under ambient conditions, suggesting that it might carry out housekeeping functions, as is the case in Arabidopsis (Lee *et al*., 2007; Myouga *et al*., 2006; Zybailov *et al*., 2009). However, in our Chlamydomonas *clpb3* mutants we found no obvious chloroplast development phenotype (Fig. S5a) and no PSII phenotype (Fig. S5b) under ambient conditions. A mild growth phenotype was observed especially under photoautotrophic conditions in mutant *clpb3-1* (Fig. 7). Obvious phenotypes under ambient conditions were neither observed in tomato *clpb3* antisense lines (Yang *et al*., 2006) nor in *Synechococcus sp. clpb3* knock-out lines (Eriksson and Clarke, 1996). Both Chlamydomonas *clpb3* mutants accumulate CLPB3 to ∼20% of wild-type levels (Figs. 2 and 3). While the 20% residual CLPB3 in mutant *clpb3-2* represent wild-type protein, this residually accumulating CLPB3 in mutant *clpb3-1* is truncated at its C-terminus (Figs. 2 and 3). If the mutagenesis cassette is indeed flanked by random DNA at its 3’ end, as indicated by our genotyping efforts (Fig. S3), the truncation comprises a stretch of ∼20 amino acids that is highly conserved among ClpB3 family members, as well as a non-conserved stretch of 52 amino acids. These sequences are most likely replaced by some junk sequence until a random stop codon is encountered. While in *E. coli* ClpB the C-terminal domain has been shown to be required for oligomer stability (Barnett *et al*., 2000; Barnett and Zolkiewski, 2002), we can only say it is important for CLPB3’s stability and functionality for the following reasons: first, the truncation obviously leads to a reduced accumulation of the protein (Fig. 2). Second, truncated CLPB3 appears to form aggregates already under ambient conditions to which complementing wild-type CLPB3 is attracted (Fig. 5). Third, truncated CLPB3 massively accumulates in aggregates during heat stress (Fig. 6). Fourth, the *clpb3-1* mutant is much more impaired in its ability to resolve heat-induced protein aggregates than the *clpb3-2* mutant, albeit both accumulate similar levels of residual CLPB3 (Figs. 2 and 6). Fifth, the *clpb3-1* mutant is more thermosensitive than the *clpb3-2* mutant (Fig. 7). Nevertheless, we did observe some CLPB3 monomers to accumulate during heat stress and recovery in *clpb3-1*. CLPB3 monomers accumulate strongly in the wild type and may indicate oligomer dynamics that are known to be important for ClpB activity (Mogk *et al*., 2015). Hence, a chloroplast development phenotype in the *clpb3-1* mutant might be concealed by some residual activity of the truncated protein, as it might be concealed by residual CLPB3 in tomato antisense lines. Clean knockout lines created e.g. by CRISPR-Cas9 will be required to solve this question in future work.

## Abbreviations

CLP: casein lytic proteinase;
HSP22: heat shock protein 22;
TIG: trigger factor;
PRPL: plastid ribosomal protein of the 50S subunit

## Supplementary Data

**Fig. S1**. Alignment of amino acid sequences of CLPB proteins from *E*.*coli* and chloroplasts.

**Fig. S2**. Production of recombinant CLPB3 in *E*.*coli*.

**Fig. S3**. Analysis of the CIB1 integration sites in the *CLPB3* gene by PCR.

**Fig. S4**. Putative heat shock elements (HSEs) in the CLPB3 promoter.

**Fig. S5**. Chlamydomonas *clpb3* mutants display no obvious phenotype regarding chloroplast development and PSII activity.

**Fig. S6**. Screening for complemented *clpb3* mutant lines.

**Table S1**. Primers used for cloning and genotyping. Lower case letters indicate nucleotides differing from the template.

### Acknowledgements

This work was supported by the Deutsche Forschungsgemeinschaft (TRR175, project C02) and the Forschungsprofil BioComp.

## Author contributions

E.K. and J.N. performed all experiments, assisted by M.M. D.S. took the immunofluorescence images. M.S. conceived and supervised the project and wrote the paper with contributions from all other authors.

## Data availability statement

All data supporting the findings of this study are available within the paper and within its supplementary materials published online.

